# Winners in good times and bad times: Aerobic anoxygenic phototrophic bacteria profit from photoheterotrophy under carbon-rich and poor conditions

**DOI:** 10.1101/2023.12.21.572764

**Authors:** Kasia Piwosz, Cristian Villena-Alemany, Joanna Całkiewicz, Izabela Mujakić, Vít Náhlík, Jason Dean, Michal Koblížek

## Abstract

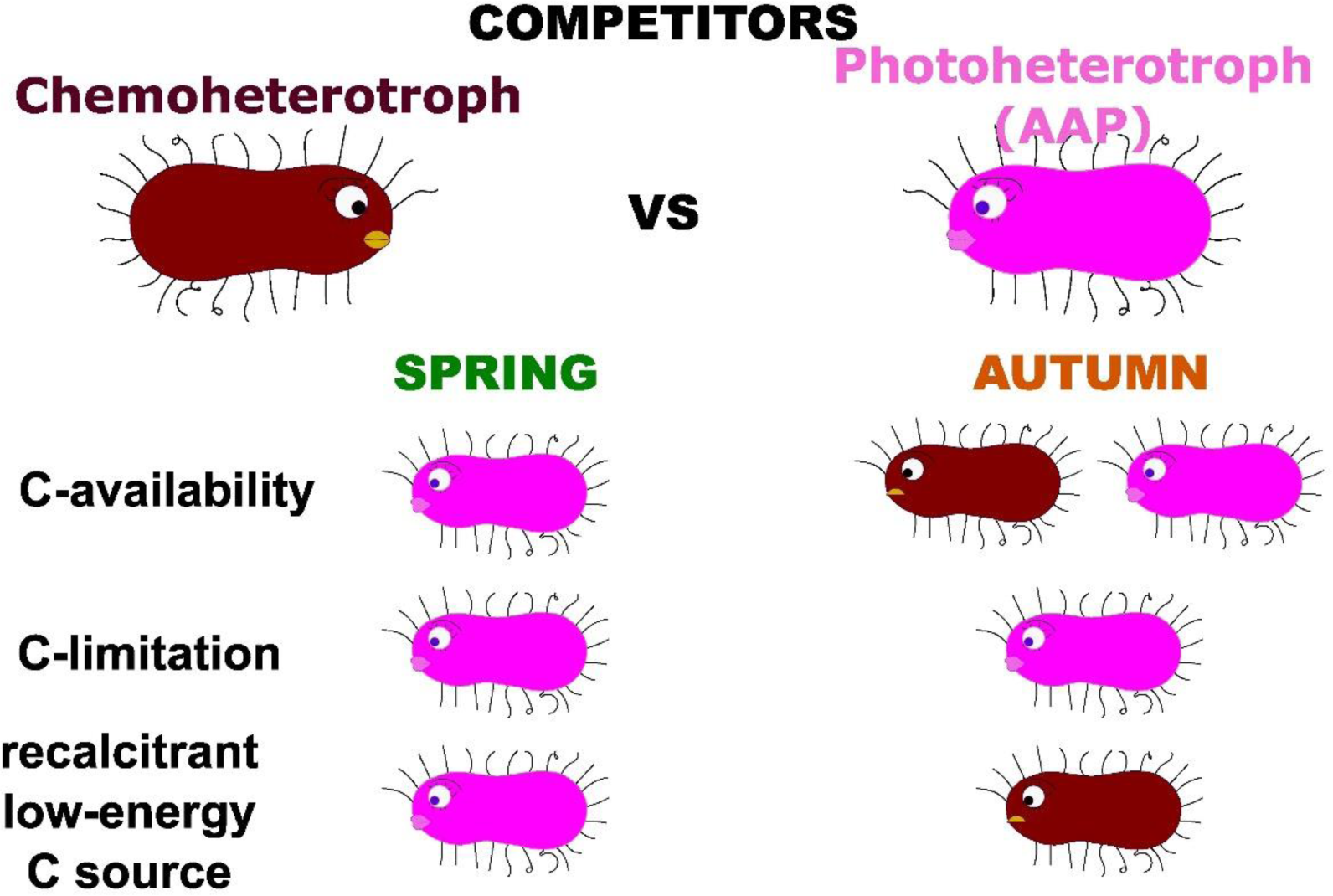

Aerobic Anoxygenic Phototrophic (AAP) bacteria are an important component of freshwater bacterioplankton. They can support their heterotrophic metabolism with energy from light, and by that enhance their growth efficiency. Based on results from cultures, it was hypothesized that photoheterotrophy provides an advantage under carbon limitation and facilitates access to recalcitrant or low-energy carbon sources. However, verification of these hypotheses for natural AAP communities has been lacking. Here, we conducted whole community manipulation experiments and compared the growth of AAP bacteria under carbon limited and with recalcitrant or low-energy carbon sources under dark and light conditions to elucidate how they profit from photoheterotrophy. We found that it depends on the season. In spring, AAP bacteria induce photoheterotrophic metabolism under carbon limitation but they outperform heterotrophic bacteria when carbon is available. This effect seems to be driven by physiological responses rather than changes at the community level. In autumn photoheterotrophy is less beneficial. In both seasons, AAP bacteria responded negatively to recalcitrant or low-energy carbon sources in light. This unexpected observation may have ecosystem-level consequences as lake browning continues. In general, our findings contribute to the understanding of the dynamics of AAP bacteria observed in pelagic environments.

## Introduction

Photoheterotrophic bacteria are an abundant part of bacterioplankton. These organisms depend on organic matter for their growth but they can supplement their energy requirements by light. One of the main photoheterotrophic groups is Aerobic Anoxygenic Phototrophic (AAP) bacteria, which harvest light by bacteriochlorophyll and carotenoid molecules bound to photosynthetic complexes to produce ATP via cyclic photophosphorylation (Okamura *et al*., 1986, Yurkov & Beatty, 1998). Therefore, when illuminated, AAP bacteria can minimize the use of organic substrates for respiration and utilize them instead for building biomass (Hauruseu & Koblížek, 2012, Piwosz *et al*., 2018b, Koblížek *et al*., 2020).

AAP bacteria were discovered in coastal marine waters (Shiba *et al*., 1979, Shiba *et al*., 1991). Later, they were also found to be common in the open ocean (Kolber *et al*., 2001), where they typically represent 1-10% of total bacteria (reviewed in Koblížek 2015). Initially, it was hypothesized that the photoheterotrophy represents an advantage in nutrient-poor oceans, which seems to be correct for rhodopsin-containing photoheterotrophs, but AAP bacteria prefer more productive coastal areas (Gómez-Consarnau *et al*., 2019, Vrdoljak Tomaš *et al*., 2019).

AAP bacteria are also commonly found in freshwater lakes, representing from < 1 to > 30% of total bacteria, with strong seasonality (Yurkov & Gorlenko, 1990, Masin *et al*., 2008, Čuperová *et al*., 2013, Lew *et al*., 2015, Kolářová *et al*., 2019). AAP cells are on average larger than heterotrophic bacteria, thus they contribute more to the total bacterial biomass than their abundance alone would indicate (Fauteux *et al*., 2015). They are also more active and have higher growth rates (Cepáková *et al*., 2016; Garcia-Chaves *et al*., 2016) thus they play an important role in the microbial food webs both as consumers of phytoplankton-derived dissolved organic matter (Piwosz *et al*., 2020) and as a food source for bacterivores (Ruiz-González *et al*., 2020). Moreover, infra-red (IR) light exposure (selectively absorbed by AAP bacteria) reduced the total microbial respiration and enhanced total microbial production (Piwosz *et al*., 2022). The extent of the effect varied over the season. The driving factors likely included seasonal succession of phytoplankton, which may significantly affected the spectrum and availability of organic substrates in the lake, and large seasonal changes in AAP community composition, which may have affected its functioning (Villena-Alemany *et al*., 2023a). However, how the additional energy from light is utilized to provide an advantage for AAP bacteria in the environment remains unknown. In addition to the aforementioned hypothesis on survival in the oligotrophic environment, it was also suggested that photoheterotrophy may help AAP bacteria to access low energetic and recalcitrant carbon sources (Salka *et al*., 2014, Koblížek, 2015).

Here, we conducted the whole microbial community manipulation experiment from a freshwater lake to test hypotheses that (i) the additional energy from light provides AAP bacteria additional energy under carbon limiting conditions, and (ii) the additional energy from light allows AAP bacteria to access low energetic or recalcitrant carbon sources. The incubations were done in the dark and in the near-infrared (IR) light (λ>800 nm), which selectively excited the infra-red *in vivo* absorption band of bacteriochlorophyll (Kasalický *et al*., 2018, Kopejtka *et al*., 2020). To all other microorganisms, both conditions were perceived as dark. We followed the bulk growth rates of heterotrophic and AAP bacteria in dark vs IR-light in the conditions of (i) organic carbon limitation, (ii) low energetic or recalcitrant organic carbon source, and (iii) natural organic carbon availability (control). We expected that in the conditions that favour photoheterotrophic metabolism, AAP bacteria would grow faster in the IR light and as a result they would increase their contribution to the total bacterial abundance Moreover, to account for the metabolic differences between different AAP phylotypes, we also followed the changes in their community.

## Material and methods

### Setting up the experiment

We conducted two experiments: in June and October 2018, to account for seasonal differences in bacterial and AAP community composition (Villena-Alemany *et al*., 2023a). Water was collected from 0.5 m depth of the meso-oligotrophic freshwater lake Cep (48°55’29.7“N, 14°53’12.5”E) using a Ruttner Water Sampler (model 11.003KC Denmark AS) on the 20^th^ of June 2018 and the 1^st^ of October 2018. It was transported to the laboratory within half an hour in a closed plastic container, which was pre-rinsed three times with the sampled water and stored in a cool box.

Two different treatments were prepared: carbon limitation (C-limited) and recalcitrant organic carbon source (lignin) in June and low-energy organic carbon source (acetate) in October (Fig. 1). For the C-limited treatment, the untreated water from the lake was diluted at a 1:4 ratio with unamended sterile inorganic basal (IBM) medium (Hahn *et al*., 2003). For the lignin/acetate treatments, the untreated water from the lake was diluted 1:4 with a sterile IBM medium containing 2.5 mg L^-1^ of dissolved lignin (in the June experiment) or 3.0 mg L^-1^ of acetate (in October). The media were prepared during the week before the experiment. They were filtered through a 0.2 µm filter and autoclaved. As a control, we used the untreated water from the lake diluted 1:4 with sterile filtered lake water that was collected three days before the experiment. It was sequentially prefiltered through a 20 µm plankton net, 0.2 µm filter, and 1-litre Stericup® Filter Units with a membrane pore size of 0.1 µm (Millipore, Merck) and kept in the dark at 4°C until the experiment. The dilution allowed for an increase in the carbon availability for bacteria and to reduce the grazing pressure from protistan grazers.Each treatment was divided into six 2-litre portions that were incubated in the dark or IR light at *in situ* temperature (21°C in June and 17°C in October) in triplicates. Samples were taken every 12 hours in June and every 24 hours in October.

**Figure 1.**
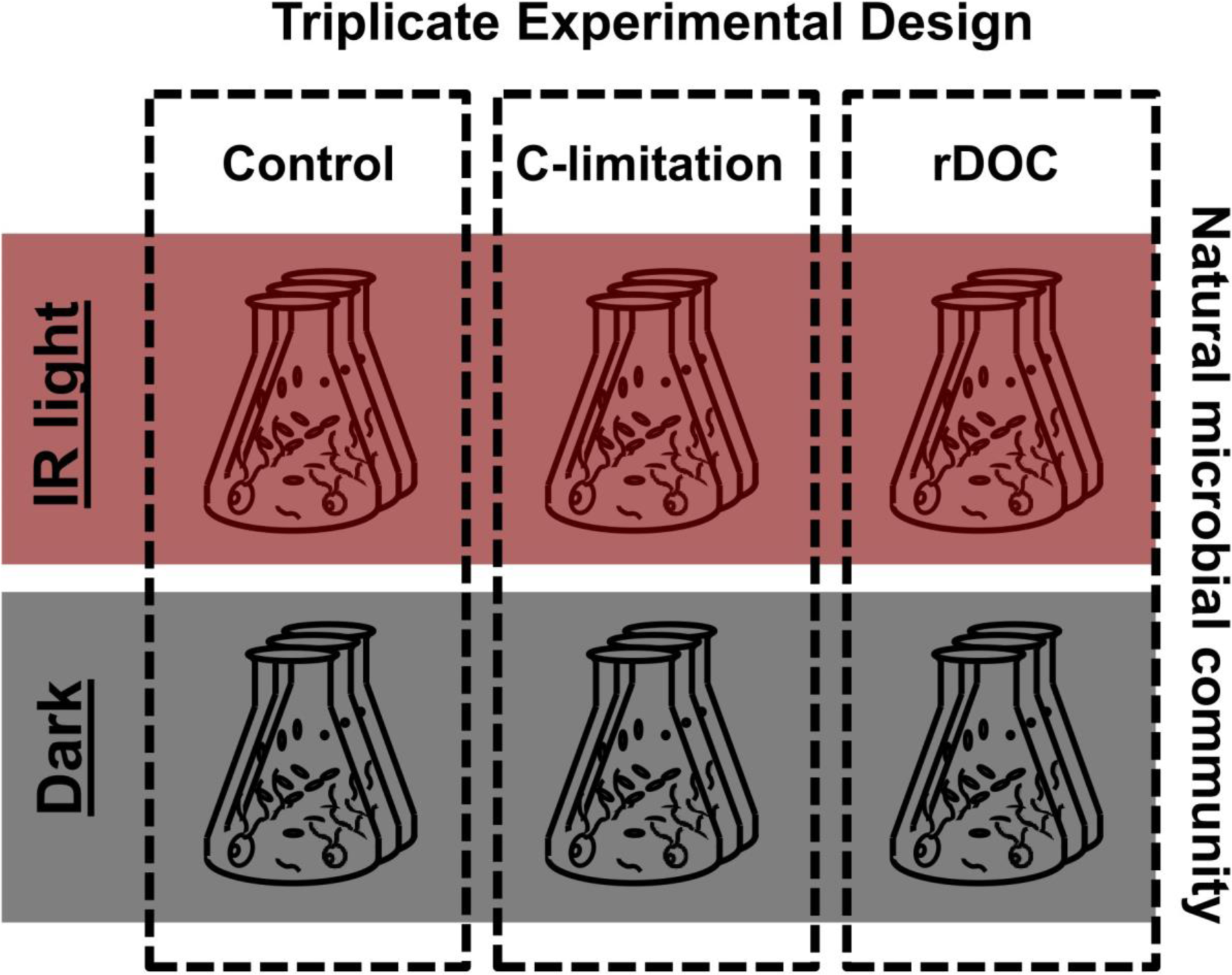
Experimental design. Natural microbial communities were diluted 1:4 with sterile filtered lake water (Control), sterile inorganic medium (C-limited) or sterile inorganic medium containing lignin (in June) or acetate (in October) as carbon source, and incubated in the dark or under infrared illumination (IR light). All treatments were performed in triplicate.

### Bacterial and AAP abundance

Samples of 50 mL were fixed with buffered, sterile-filtered paraformaldehyde (Penta, Prague, Czechia) to a final concentration of 1%, and 0.5 mL was filtered onto white polycarbonate filters (pore size 0.2 µm, Nucleopore, Whatman, UK). Cells were stained with 4’,6-diamidino-2-phenylindole at a concentration of 1 mg L^-1^ (Coleman, 1980). Total and AAP bacterial abundances were determined using an epifluorescence Zeiss Axio Imager.D2 microscope (Cottrell *et al*., 2006, Villena-Alemany *et al*., 2023a). The abundance of heterotrophic bacteria was calculated as the difference between the total bacteria and AAP bacteria.

### DNA extraction, amplicon preparation and sequencing

About 350 mL of water was filtered onto sterile 0.2 µm Nucleopore Track-Etch Membrane filters (Whatman, Maidstone, United Kingdom). Filters were placed inside sterile cryogenic vials containing 0.55 g of sterile zirconium beads, flash-frozen in liquid nitrogen and stored at −80 °C until DNA extraction (< 6 months). Total nucleic acids were chemically extracted according to Griffiths et al. (2000) with modifications (Nercessian et al. 2005). Extracted DNA was re-suspended in 35 µl of DNase and RNase-free water (MP Biomedicals, Solon, OH, USA) and stored at −20 °C. The concentration and quality of the extracts were checked using a NanoDrop (Thermo Fisher Scientific). A pure culture of *Dinoroseobacter shibae* was used as a control for cross-contamination between the samples.

Amplicons for the *puf*M gene (a marker gene for AAP bacteria) were prepared using *puf*M UniF and *puf*M UniR primers (Yutin *et al*., 2005). The PCR conditions were as follows: initial denaturation for 3 min at 98°C, 30 cycles of 98°C for 10 s, 52°C for 30 s, 72°C for 30 s, and final elongation at 72°C for 5 min. PCR was performed in 20 μL triplicate reactions using Phusion™ High-Fidelity PCR MasterMix (ThermoFisher Scientific, USA).

The triplicate reactions for each sample were pooled and purified from the gel using the Wizzard SV Gel and PCR clean system (Promega) and quantified using the Qubit dsDNA HS assay (ThermoFisher Scientific). Amplicons were pooled in equimolar amounts and sequenced on the Illumina MiSeq (2×250 bp) platform at the Genomic Service of the Universitat Pompeu Fabra (Barcelona, Spain).

### Analysis of amplicon data

Reads were quality-checked using FastQC v0.11.7 (Babraham Bioinformatics, Cambridge, UK). Primer sequences were trimmed in Cutadapt v1.16 (Martin, 2011). Subsequent analyses were done in the R/Bioconductor environment using the DADA2 package (version 1.12.1) (Callahan *et al*., 2016). *puf*M sequences were processed and assigned using reference database and methods described in Villena-Alemany *et al*. (2023b). The contamination in the *D. shibae* culture was about 1%. To remove this contamination, the final ASVs tables consisted of ASVs with the sum of reads in all samples > 10 and present in at least two replicates in a treatment at a given time point, or with the sum of reads in all samples > 10 and present in at least three time points in a given replicate in a treatment (Piwosz *et al*., 2018a).

### Statistical analysis

Growth rates were calculated as linear fit coefficients on abundance data transformed with natural logarithm. Differences between incubation in the dark and IR treatment at the end of the experiment were tested with the Welsh t-test. The p-value was adjusted for multiple tests using Bonferroni correction, and the significance of the results was assumed for p<0.01. The distribution of the data was tested with the Shapiro-Wilk test. The changes in ASVs’ reads abundance between control and C-limited, and control and Lignin/Acetate treatments in the IR light at the end of the experiments were tested using the DESeq function (test=“Wald”, fitType=“parametric”) from DESeq2 package (version 1.36.0). All analyses were done in Rstudio for Windows (version 2023.03.1+446; R version 4.2.0 (The R Core Team, 2015).

### Data availability

The sequences were deposited in the EMBL database under Biosamples ERS17465032-ERS17465210 and ERS17468627 in the BioProject PRJEB71033.Count data are available as Supplementary Table S1.

## Results

### June experiment

Heterotrophic bacteria grew the fastest in the control treatment (IR light: 0.31±0.03 d^-1^, dark: 0.25±0.05 d^-1^, p=0.57), where they almost doubled within 56 h (Fig. 2a). The growth rate was slower in the C-limited (IR light: 0.26±0.13 d^-1^, dark: 0.16±0.03 d^-1^, p=0.37), while in the Lignin treatment they almost did not grow (IR light: −0.05±0.06 d^-1^, dark: 0.12±0.02 d^-1^, p=0.016).

**Figure 2.**
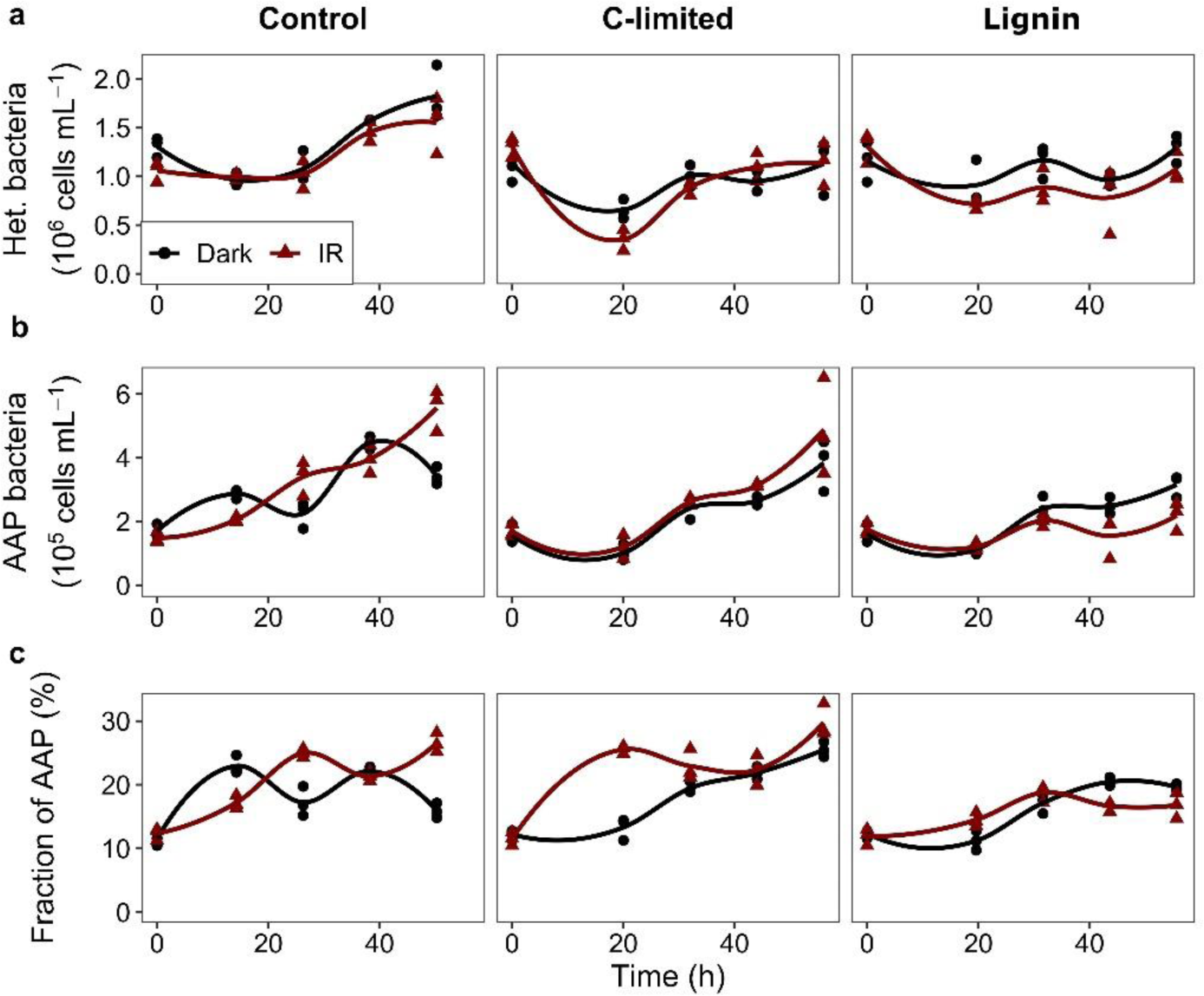
Abundance of heterotrophic bacteria (a); AAP bacteria (b) and contribution of AAP bacteria to total bacterial numbers (c) in the June experiment. Values for each triplicate are shown as points, and the line was fitted locally using the loess function from the ggplot2 package in R.

The effect of IR light on AAP bacteria was evident in the control treatment: their growth rate was almost twice as fast in the IR light than in the dark (0.66±0.02 d^-1^ vs 0.37±0.07 d^-1^, respectively, p < 0.006). This resulted in a higher abundance (IR light: 5.57±0.66×10^5^ cells ml^-1^, dark: 3.43±0.27×10^5^ cells ml^-1^, p<0.009) and contribution of AAP bacteria (IR light: 26.6±1.5%, dark: 15.9±1.2%, p<0.0005) in the IR light at the end of the experiment (Fig. 2b, c). A steady growth of AAP bacteria was also observed in the C-limited treatment in both dark and IR light conditions, but the difference in growth rate (0.55±0.06 d^-1^ in IR light and 0.55±0.04 d^-1^ in dark, p=0.56), abundance (IR light: 4.07±0.55 d^-1^ ×10^5^ cells ml^-1^, dark: 3.84±0.81×10^5^ cells ml^-1^, p>0.35) and contribution (IR light: 29.8±2.6%, dark: 25.5±1.2%, p<0.043) was insignificant. Furthermore, the differences in growth rate, abundance and contribution of AAP bacteria between the control and C-limited treatments in the IR light were insignificant. Interestingly, the growth of AAP bacteria was slower in IR light in the Lignin treatment (IR light: 0.12±0.09 d^-1^, dark: 0.43±0.02 d^-1^, p=0.01), which resulted in their slightly higher abundances (IR light: 2.19±0.45×10^5^ cells ml^-1^, dark: 3.16±0.36×10^5^ cells ml^-1^, p>0.02) and contributions (IR light: 16.8±2.0%, dark: 19.6±0.5%, p=0.13) in the dark. The growth of AAP bacteria in the IR light was significantly slower in the Lignin treatment compared to the control (p=0.003), resulting in their lower abundances (p=0.0015) and contribution (p=0.0017) at the end of the experiment. AAP bacteria grew more than twice as fast as heterotrophic bacteria in the IR light in all treatments, indicating that they profited from photoheterotrophy under all of these conditions.

The changes in AAP community composition were minor and only several ASVs significantly altered their relative abundance in C-limited and Lignin treatments compared to the control at the end of the experiment in the dark but not in the IR light (Fig 3, Supplementary Table S2). C-limitation induced a relative increase of *Novosphingobium* and *Methylobacterium* (Alphaproteobacteria), and *Limnohabitans* (Gammaproteobacteria) compared to the control treatment. Members of the genus *Limnohabitans* were also affected by Lignin treatment, with different ASVs either profiting or losing in these conditions (Fig. 3c).

**Figure 3.**
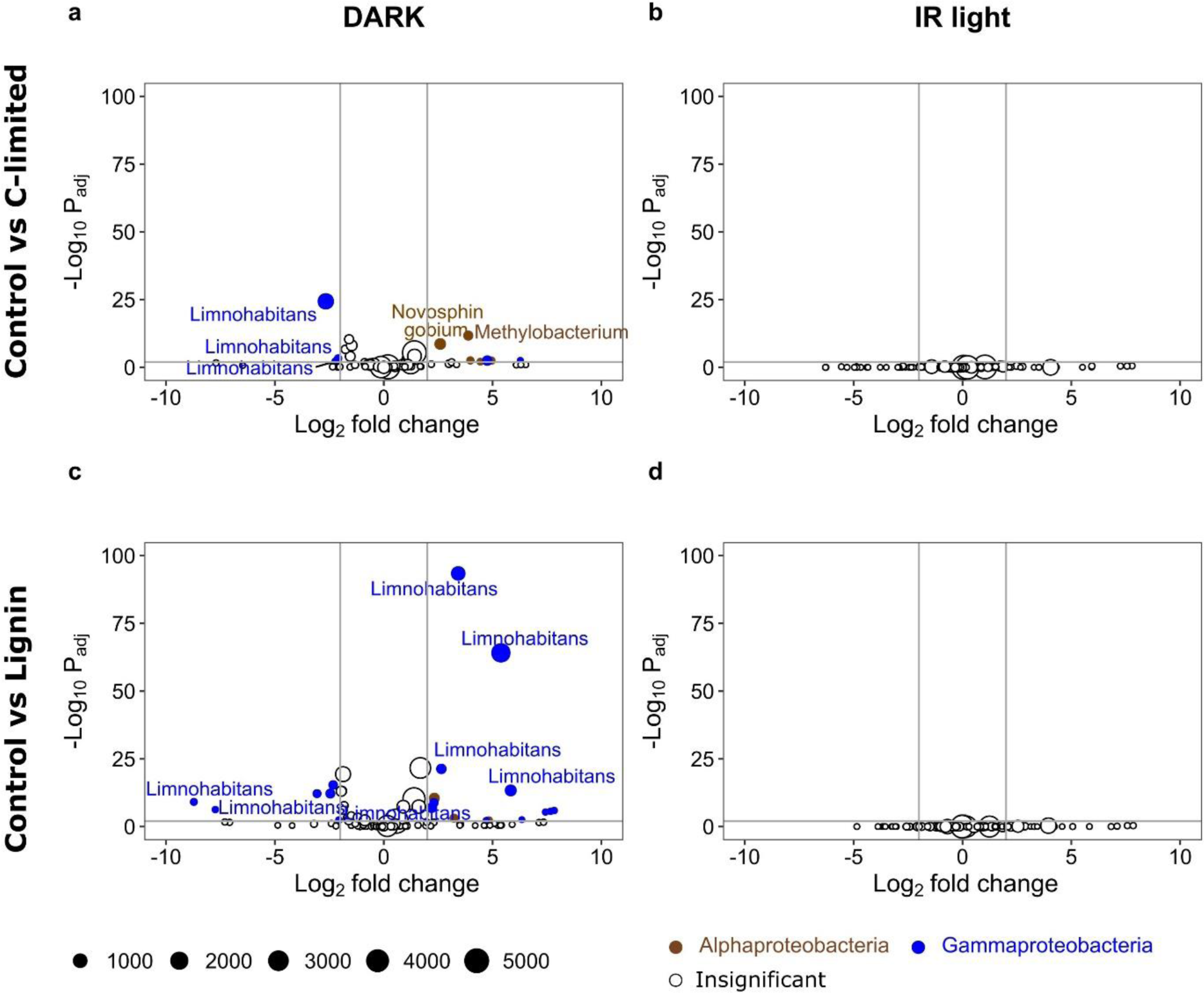
Vulcan plots showing the ASVs with significantly different (adjusted p-value < 0.01, Log2 fold change > |2|) relative abundance at the end of the experiment between (a-b) control and C-limited treatments, and (c-d) control and Lignin treatments in dark (a, c) and IR light (b, d) in June experiment. Negative Log2 fold change value (x axes) indicates that the read count of an ASV was lower in the experimental treatment than in the control, and the positive that it was higher. Vertical grey lines show Log2 fold change values of −2 and 2, horizontal grey lines show significance level (adjusted p-value < 0.01). Bubble size corresponds to the mean number of reads in both compared treatments, colours show the Class affiliation for significant ASVs (brown – Alphaproteobacteria, blue – Gammaproteobacteria, white – insignificant).

### October experiment

The growth patterns in the October experiment were different than in June. Heterotrophic bacteria grew fastest in the C-limited treatment regardless of the light conditions (IR light: 0.16±0.02 d^-1^, dark: 0.17±0.01 d^-1^, p=0.71), reaching similar abundance at the end of the experiment (Fig. 4a). Their growth rate was twice as fast in the IR light as in the dark in the control treatment (IR light: 0.12±0.04 d^-1^, dark: 0.06±0.03 d^-1^, p=0.06), but the abundance at the end of the experiment was similar (IR light: 1.59±0.27×10^6^ cells ml^-1^, dark: 1.35±0.09×10^6^ cells ml^-1^, p=0.13). In contrast, in the Acetate treatment, heterotrophic bacteria grew only in the dark (growth rate IR light: −0.04±0.05 d^-1^, dark: 0.16±0.01 d^-1^, p=0.01), reaching much higher abundances at the end of the experiment (IR light: 1.30±0.15×10^6^ cells ml^-1^, dark: 2.29±0.36×10^6^ cells ml^-1^, p=0.01).

**Figure 4.**
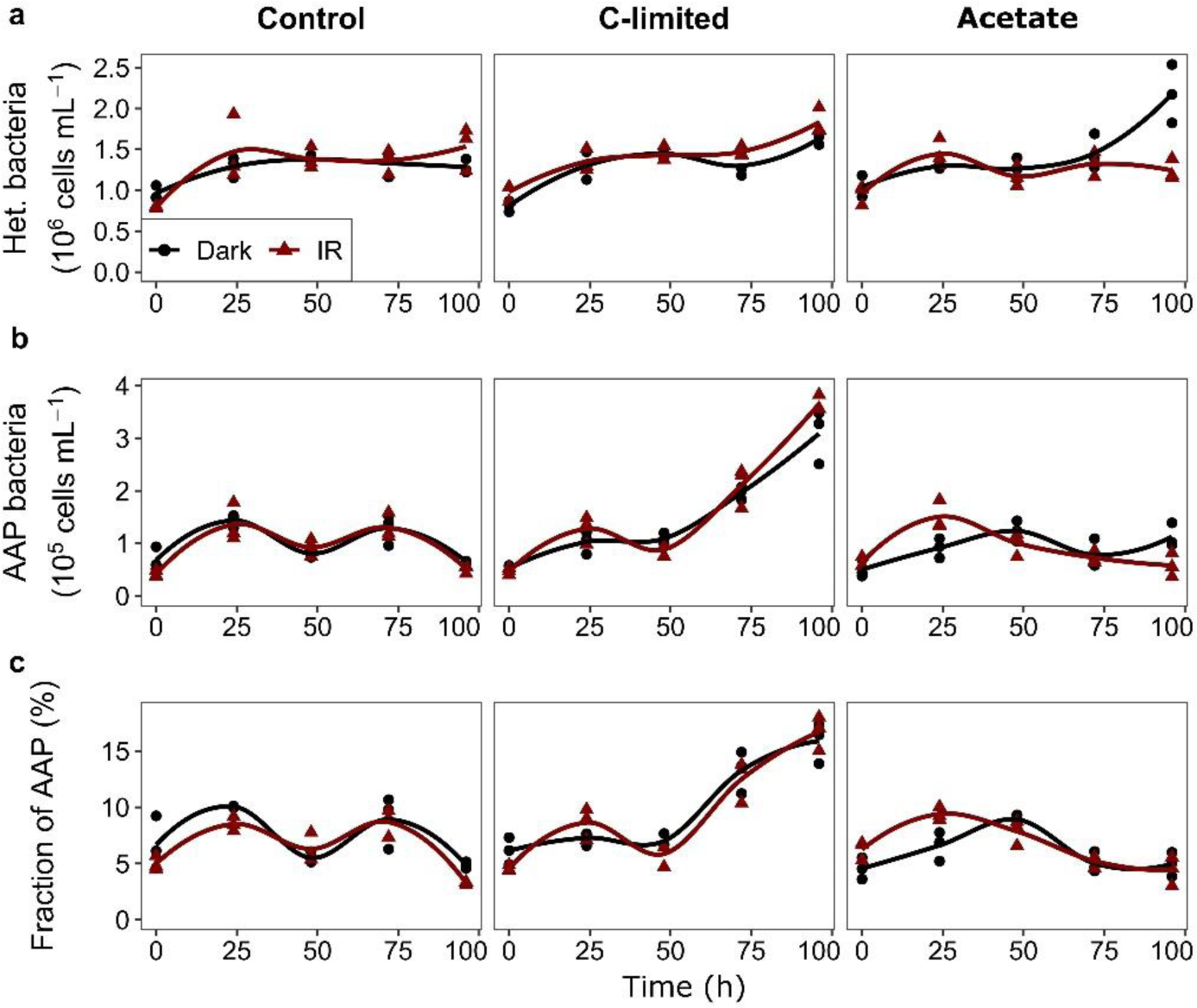
Abundance of heterotrophic bacteria (a); AAP bacteria (b) and contribution of AAP bacteria to total bacterial numbers (c) in the October experiment. Values for each triplicate are shown as points, and the line was fitted locally using loess from the ggplot2 package in R.

The growth rate of AAP bacteria was close to 0 in the control treatment, both in the IR light (0.04±0.05 d^-1^) and dark (−0.01±0.03 d^-1^, p=0.11), and their abundance and contribution to total bacterial abundance did not change (Fig. 4b, c). In contrast, they grew rapidly in the C-limited treatment in both dark and light conditions (growth rate IR light: 0.46±0.05 d^-1^, dark: 0.42±0.07 d^-1^, p=0.02, Fig. 4b) and their contribution to the total bacterial abundance tripled (IR light: 16.7±1.5%, dark: 15.9±1.8%, p<0.02, Fig. 4c). Their growth rate, final abundance and contribution were significantly higher in the C-limited treatment than in the control (p<0.003). Finally, in the Acetate treatment, AAP bacteria grew in the dark (0.15±0.10 d^-1^) but decreased in the IR light (−0.37±0.02 d^-1^, p=0.006), resulting in twice lower abundance in the IR light at the end of the experiment (IR light: 0.58±0.22×10^5^ cells ml^-1^, dark: 1.13±0.22×10^5^ cells ml^-1^, p=0.02). However, even though the growth rate was significantly lower in the Acetate than in the control (p=0.002), their abundance and contribution at the end of the experiment did not differ between these treatments (p>0.6).

The growth rate of heterotrophic bacteria compared to AAP bacteria in the IR light did not differ in the control treatment (p=0.94), was significantly lower in the C-limited treatment in IR light (p= 0.0023), and was significantly higher in the IR light in the Acetate treatment (p=0.003).

As observed in June, only several ASVs significantly changed their relative abundance during the experiment both in the dark and IR light (Supplementary Table S3). In the control treatment, *Limnohabitans* and *Hydrogenophaga* increased, while Methylobacteriaceae and Gemmatimonadaceae decreased, especially in the IR treatment. In the C-limited and Acetate treatments, the ASVs that increased were affiliated mainly with *Hydrogenophaga*, while those that decreased included Methylobacteriaceae, Gemmatimonadaceae, Pseudomonadales UBA5518 and other Burkholderiaceae (Supplementary Table S3). The number of ASVs that showed significantly different relative abundance at the end of the experiment between the control and C-limited or Acetate treatments was lower than that observed within the treatments between T0 and T end (Supplementary Table S3). *Hydrogenophaga* increased in the C-limited and Acetate treatments compared to the control both in the dark and IR light, while *Limnohabitans* and *Sandarakinorhabdus* decreased but only in the dark (Fig. 5).

**Figure 5.**
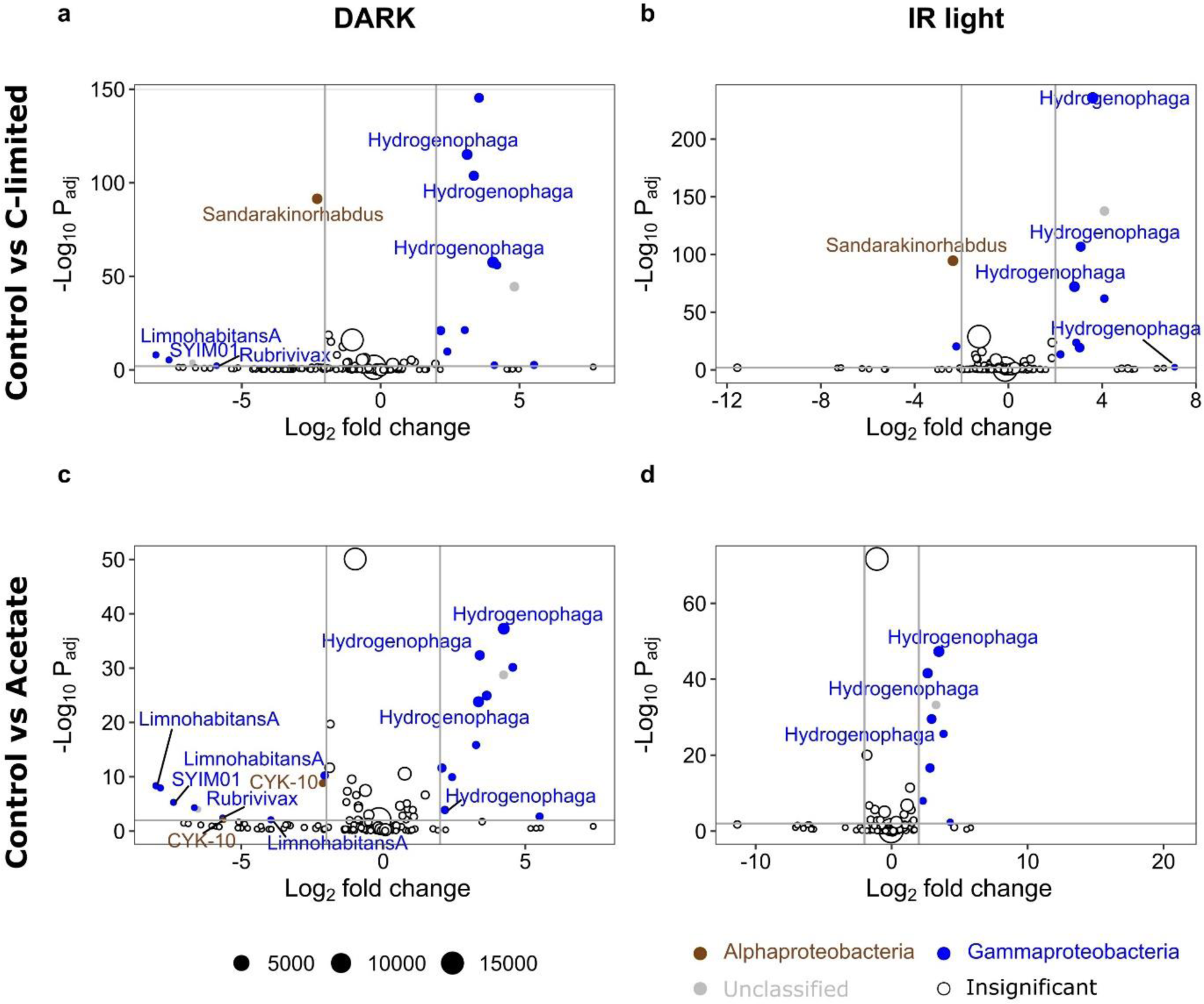
Vulcan plots showing the ASV with significantly different (adjusted p-value < 0.01, Log2 fold change > |2|) relative abundance at the end of the experiments between (a-b) control and C-limited treatments, and (c-d) control and Acetate treatments in dark (a,c) and IR light (b, d) in the October experiment. Negative Log2 fold change value (x axes) indicates that the read count of an ASV was lower in the experimental treatment than in the control, and the positive that it was higher. Vertical grey lines show Log2 fold change values of −2 and 2, horizontal grey lines show significance level (adjusted p-value < 0.01). Bubble size corresponds to the mean number of reads for both compared treatments, colours show the Class affiliation for significant ASVs (brown – Alphaproteobacteria, blue – Gammaproteobacteria, grey – unclassified, white – insignificant).

## Discussion

When AAP bacteria were discovered to be abundant in the Northeast Pacific, it was assumed that their ability to use light to produce ATP to support their heterotrophic metabolism helps them to survive in the oligotrophic environment of the ocean (Kolber *et al*., 2001). However, later observations of their phenology and distribution indicated otherwise: higher concentrations were observed in more eutrophic coastal waters during or shortly after the phytoplankton bloom (Auladell *et al*., 2019, Vrdoljak Tomaš *et al*., 2019). A similar pattern emerged from freshwater studies (Kolářová *et al*., 2019, Villena-Alemany *et al*., 2023b), questioning the initial assumption. It was also suggested that additional energy from light can facilitate access to recalcitrant complex organic polymers or low-energy carbon sources, such as lignin or acetate (Koblížek, 2015). However, experimental support for these statements comes mostly from experiments with cultured species (Kuzyk *et al*., 2023), and the question of how AAPs utilize their photoheterotrophy in natural environments remains open.

The key finding of this study is that the profit from photoheterotrophic nutrition differed between the seasons. The positive response of AAP bacteria to IR light only in the control in the June experiment (Fig. 2b, c) suggests that additional energy from light allows them to successfully compete with heterotrophic bacteria at the time of surplus DOC availability in spring. This result contradicts the initial hypotheses (Kolber *et al*., 2001), but agrees with the fact that many freshwater AAP genera (*e.g. Limnohabitans*) are considered copiotrophs (Chiriac *et al*., 2023, Park *et al*., 2023). Furthermore, in June AAP bacteria grew faster than the heterotrophic bacteria in all treatments, providing experimental support for the recently suggested inclusion of AAP bacteria in the Phytoplankton Ecology Group (PEG) model as super competitors profiting from spring phytoplankton bloom (Villena-Alemany *et al*., 2023b). However, the negligible growth of AAP bacteria in the control treatment in the October experiment indicates that this advantage is irrelevant in autumn (Fig. 4b, c) when the AAP community may represent different gene repertoires and thus distinct functionalities(Villena-Alemany *et al*., 2023b). In general, AAP bacteria seem to be less active and abundant in autumn than in spring (Kolářová *et al*., 2019, Piwosz *et al*., 2022, Villena-Alemany *et al*., 2023a).

The growth of AAP bacteria in the C-limited treatment was similar in the IR light and the dark in both seasons (Fig. 2 and 4), indicating that they did not profit from photoheterotrophy in such conditions. However, this was contradicted by the observation that they grew much faster than heterotrophic bacteria, increasing their contribution to the total bacterial community up to threefold. This incongruence suggests a dual role of photoheterotrophic metabolism for AAP bacteria: On one hand, carbon limitation triggers the production of BChl-*a* to aid their survival (Kolber *et al*., 2001, Koblížek, 2015, Kopejtka *et al*., 2020, Kuzyk *et al*., 2023). On the other, photoheterotrophy seems to be the most advantageous under carbon availability, when a more efficient metabolism allows them to grow faster than heterotrophic bacteria, as discussed above.

An unexpected result was the negative response of the AAP bacteria to the Lignin treatment in June and the Acetate treatment in October (Fig. 2, 3). This indicates that both complex polymers (lignin) and low-energy monomers (acetate) are disadvantageous for freshwater AAP bacteria. This negative relationship, for which the mechanism remains yet unknown, may have serious consequences for lake functioning. Currently, many temperate lakes in the Northern Hemisphere are affected by browning, resulting from the increase in terrestrial dissolved organic matter (DOM) (Williamson *et al*., 2015) which is often recalcitrant for bacteria (Kritzberg *et al*., 2004). Browning is predicted to continue as the atmospheric acid deposition decreases and due to climate change (Meyer-Jacob *et al*., 2019), affecting pelagic food webs (Williamson *et al*., 2015) and increasing CO_2_ flux to the atmosphere (Kritzberg *et al*., 2020). While some AAP bacteria, such as *Sphingomonas* sp. strain FukuSWIS1 from the acidic lake Grosse Fuchskuhle (Salka *et al*., 2014), seem to be adapted to conditions prevailing in humic and brown lakes, our results indicate that overall AAP bacteria may be negatively impacted by recalcitrant or low energy carbon sources. This effect may be more pronounced in spring when terrestrial DOM inputs, potentially higher due to river runoff, could hamper their photoheterotrophy, thus decreasing the efficiency of carbon assimilation and lowering its availability for higher trophic levels (Piwosz *et al*., 2022, Villena-Alemany *et al*., 2023b). Further experiments employing a wider variety of recalcitrant and low-energy compounds are needed to confirm and understand this effect.

AAP community composition shows recurrent seasonal patterns in freshwater lakes (Villena-Alemany *et al*., 2023b), which are driven by changes in environmental conditions, such as temperature and DOC concentration (Villena-Alemany *et al*., 2023a). The minor changes in the AAP community composition under IR light observed here indicate that the response to the treatments could have been driven by a physiological change in specific AAP phylotypes switching to phototrophic metabolisms rather than a community-level response (Fecskeová *et al*., 2019, Piwosz *et al*., 2020). For instance, the only genus that had significantly increased its relative abundance in IR light was *Hydrogenophaga* in October (Fig. 5 b, d). Members of this genus were reported to oxidize hydrogen as an energy source (Willems *et al*., 1989), which may have interesting implications for the functional role of AAP bacteria in freshwaters. In addition, numerous ASVs affiliated with *Limnohabitans* either increased or decreased in Lignin treatment in June (Fig. 3c), which may indicate niche separation between closely related AAP species (Villena-Alemany *et al*., 2023b).

Interestingly, more ASVs changed their relative abundance under dark conditions throughout the experiment, especially in June (Figs. 3, 5). This suggests that dark incubations which are commonly used in experimental design to minimise the effect of primary producers (Šimek *et al*., 2020, Fecskeová *et al*., 2021), may exaggerate the community-level responses compared to light treatments (Piwosz *et al*., 2020). This observation aids in arguments that dark incubations provide biased insights into the activity of freshwater bacterioplankton (Piwosz *et al*., 2022).

### Conclusions

Our experimental evidence indicates that although AAP bacteria’s ability to use light as a supplementary energy source is induced under carbon limitation, they profit from photoheterotrophy when carbon is available. However, this advantage over heterotrophic bacteria depends on the season and may be more pronounced in springtime. This effect also seems to be driven by physiological responses rather than changes at the community level. These findings contribute to our understanding of the dynamics of AAP bacteria observed in pelagic environments. Finally, the observation of the negative effect of lignin and acetate on AAP bacteria opens a new research topic in their ecology, as it may have ecosystem-level consequences as lake browning continues.

## Acknowledgements

Authors Contribution (CRediT): KP: Conceptualization, Data curation, Formal Analysis, Funding acquisition, Investigation, Project administration, Resources, Supervision, Validation, Visualization, Writing – original draft; CVA: Formal Analysis, Writing – review & editing; JC: Data curation, Formal Analysis, Writing – review & editing; IM: Investigation, Writing – review & editing, VN: Investigation, Writing – review & editing; JD: Investigation, Writing – review & editing; MK: Conceptualization, Funding acquisition, Writing – review & editing.

